# Intra-granuloma accumulation and inflammatory differentiation of neutrophils underlie mycobacterial ESX-1-mediated immunopathology

**DOI:** 10.1101/2022.01.19.477031

**Authors:** Julia Lienard, Kristina Munke, Line Wulff, Clément Da Silva, Julien Vandamme, Katie Laschanzky, Thorsten Joeris, William Agace, Fredric Carlsson

**Author notes:** Corresponding author /.

## Abstract

The conserved ESX-1 type VII secretion system is a major virulence determinant of pathogenic mycobacteria, including *Mycobacterium tuberculosis* and *Mycobacterium marinum*. ESX-1 is known to interact with infected macrophages, but its potential roles in regulating other host cells and immunopathology have remained largely unexplored. Using a murine *M. marinum* infection model we identify neutrophils and Ly6C^+^MHCII^+^ monocytes as the main cellular reservoirs for the bacteria. We show that ESX-1 promotes intra-granuloma accumulation of neutrophils and that neutrophils have a previously unrecognized required role in executing ESX-1-mediated pathology. To explore if ESX-1 also regulates the function of recruited neutrophils we performed single cell RNA-seq analysis indicating that ESX-1 drives newly recruited uninfected neutrophils into an inflammatory phenotype via an extrinsic mechanism. In contrast, monocytes restricted the accumulation of neutrophils and immunopathology, demonstrating a major host-protective function for monocytes specifically by suppressing ESX-1-mediated neutrophilic inflammation. iNOS activity was required for the suppressive mechanism and we identified Ly6C^+^MHCII^+^ monocytes as the main iNOS-expressing cell type in the infected tissue. These results suggest that ESX-1 mediates immunopathology by driving neutrophil accumulation and phenotypic differentiation in the infected tissue, and they demonstrate an antagonistic interplay between monocytes and neutrophils by which monocytes suppress host-detrimental neutrophilic inflammation.

**Author Summary:** The ESX-1 type VII secretion system is required for virulence of pathogenic mycobacteria, including *Mycobacterium tuberculosis*. ESX-1 interacts with infected macrophages, but its potential roles in regulating other host cells and immunopathology have remained largely unexplored. We demonstrate that ESX-1 promotes immunopathology by driving intra-granuloma accumulation of neutrophils, which upon arrival adopt an inflammatory phenotype in an ESX-1-dependent manner. In contrast, monocytes limited the accumulation of neutrophils and neutrophil-mediated pathology via an iNOS-dependent mechanism, suggesting a major host-protective function for monocytes specifically by restricting ESX-1-mediated neutrophilic inflammation. These findings provide insight into how ESX-1 promotes disease, and they reveal an antagonistic functional relationship between monocytes and neutrophils that may regulate immunopathology not only in mycobacterial infection, but also in other infections as well as in inflammatory conditions and cancer.

## Introduction

The interplay between host-pathogen interactions and functional interactions among different types of host cells – both infected and uninfected bystanders – remains poorly understood. Still, such dynamics is likely central to infection biology, including mycobacterial pathogenesis and granuloma formation. The ESX-1 type VII secretion systems in *Mycobacterium tuberculosis* and *Mycobacterium marinum* are highly conserved [1] and interact similarly with infected macrophages to promote intracellular growth [2, 3] and to regulate cytokine output [4–6]. Studies with ESX-1-deficient mutants in both species indicate required roles for this secretory system in virulence [3, 7–10] and granuloma formation [7, 9, 10]. Attenuation of the *Mycobacterium bovis* bacillus Calmette-Guérin vaccine strain is largely explained by a loss of ESX-1 [11], further emphasizing the general importance of this secretory system in mycobacterial pathogenesis. However, our knowledge regarding the mechanisms by which ESX-1 promotes disease is focused on its interactions with infected macrophages, and little is known about how it might affect other cell types and how it promotes pathology *in vivo*. To explore these questions we employed a recently established mouse model of *M. marinum* infection in which the bacteria grow and cause granulomatous disease specifically in tail tissue, a feature caused by the low optimal growth temperature (∼32°C) of the bacteria and the cooler environment in the tail [7]. Importantly, in contrast to the murine *M. tuberculosis* infection model this model allows for longitudinal and quantitative analysis of disease progression in live animals, and it exhibits the formation of caseating granulomas with an architecture similar to those formed in human tuberculosis [7].

Neutrophils are a hallmark of mycobacterial granulomas and are associated with active disease [12] and pulmonary destruction [13] in patients. Recent studies have identified neutrophils as a cellular reservoir for *M. tuberculosis* in humans [14] and experimental models [15–18]. In mice, neutrophils exhibit CXCR2-dependent recruitment to the site of infection [19, 20], where cognate chemokine ligands are produced via an IL1-mediated process that can be inhibited by nitric oxide [19, 21]. While several studies suggest a host-detrimental role for neutrophils in mycobacterial infection [18-20, 22-24] their role remains controversial [25–28], and the host-pathogen interactions and mechanisms that regulate neutrophil recruitment and function have not been elucidated. Mononuclear phagocytes are key constituents of granulomatous lesions and represent an important cellular reservoir for *M. tuberculosis* in both humans [29] and animal models [15–18]. In the tissue, newly recruited monocytes rapidly differentiate into functionally diverse subsets [30, 31], which may harbor bacteria within a few days of their arrival [30]. *M. tuberculosis* infected monocyte-deficient CCR2^-/-^ mice exhibit exaggerated disease [32–35] and delayed CD4^+^ T cell activation [33, 34]. However, whether monocytes are able to actively suppress pathology – and if so, by what mechanism – has remained an open question.

## Results

### *M. marinum* drives neutrophilic inflammation in an ESX-1-dependent manner

C57Bl/6 mice were infected with wild type (WT) or an isogenic ESX-1-deficient mutant (ΔRD1) of *M. marinum* via tail vein injection and analyzed for development of visible lesions and bacterial growth. As expected [7], ESX-1 promoted disease development (S1A-B Fig.) and bacterial growth in the tail (S1C Fig.), and bacteria were unable to propagate in the spleen (S1C Fig.). Consistent with the localized nature of the infection, the bacterial load in tail tissue was mirrored in tail draining sciatic and inguinal lymph nodes but not in intestinal draining mesenteric lymph nodes (S1C Fig.).

Kinetic flow cytometry analysis of WT and ΔRD1 infected tail tissues demonstrated a significant role for ESX-1 in driving influx of CD45^+^ hematopoietic cells (Fig. 1A). More detailed analysis showed that ESX-1 promotes significant accumulation of CD11b^+^Ly6G^+^ neutrophils by 14 days post infection (Fig. 1B-C and S1D Fig.). The increase of neutrophils was associated with ESX-1-dependent production of CXCL1, 2 and 5 – chemokine ligands for CXCR2 [20] – in the infected tissue (S1E Fig.). CD64^+^ myeloid cell numbers were slightly elevated in an ESX-1-independent fashion while the number of CD11c^+^MHCII^+^ conventional dendritic cells remained unaffected by infection (Fig. 1B-C and S1D Fig.). Subdivision of the CD64^+^ compartment into Ly6C^hi^MHCII^-^ monocytes (Gate 1; G1), Ly6C^+^MHCII^+^ monocytes (G2), Ly6C^-^MHCII^+^ monocytes (G3), and Ly6C^-^MHCII^low^ tissue-resident macrophages (G4) showed that infection caused an increase primarily of G2 monocytes whereas G4 macrophage numbers were reduced (Fig. 1D-E and S1F Fig.). B cell cellularity remained similar throughout the infection, and CD4^+^ and CD8^+^ T lymphocytes increased modestly with a significant contribution of ESX-1 (Fig. 1F-G and S1G Fig.).

**Fig. 1.**
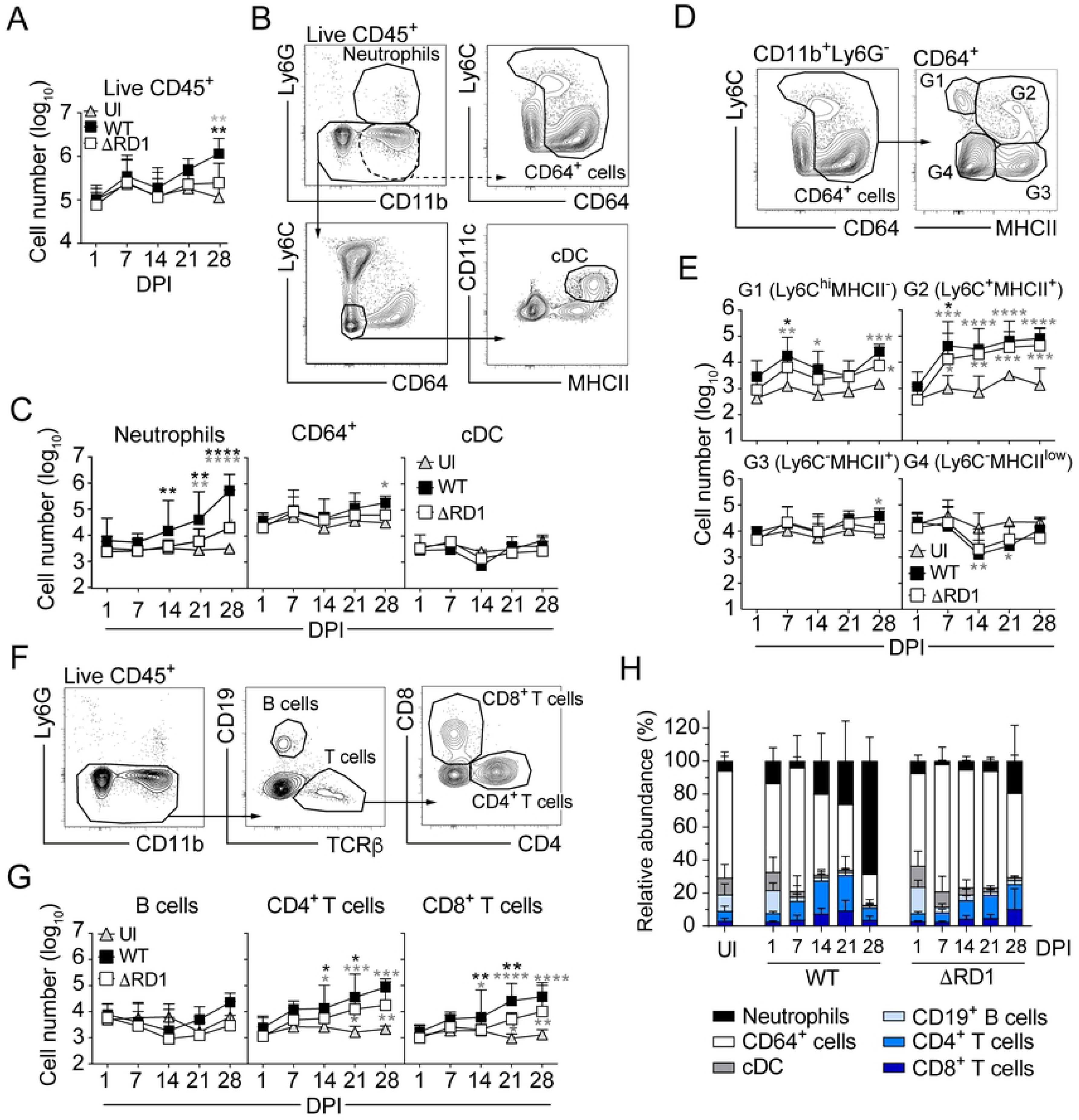
*M. marinum* drives neutrophilic inflammation in an ESX-1-dependent manner. Flow cytometry analysis of tail tissues from C57Bl/6 mice infected with 5 x 10^6^ CFUs of WT or ΔRD1 *M. marinum*, or left uninfected (UI), as indicated. **A**) Number of hematopoietic cells (CD45^+^). **B**) Gating strategy defining neutrophils (Ly6G^+^CD11b^+^), monocytes-derived cells and macrophages (CD64^+^) and conventional dendritic cells (cDC; CD64^-^MHCII^+^CD11c^+^) in an UI mouse. **C**) Quantification of data as defined in B. **D**) Subdivision of CD64^+^ cells into G1 to G4 gates based on MHCII and Ly6C expression. **E**) Number of cells within the G1 to G4 gates as defined in D. **F**) Gating strategy to define B (CD19^+^) and T cells (TCRβ^+^CD4^+^ or CD8^+^). **G**) Quantification of data as defined in F. **H**) Relative abundance of each cell population among the 6 defined populations in B and F. (**A**-**H**) Results (mean + SD, *n* = 7-9 infected and 2-3 uninfected mice) from two independent experiments. Black stars; comparison between WT and ΔRD1 infected mice. Grey stars; comparison between infected and UI mice. 2-way ANOVA. *p<0.05, **p<0.01, ***p<0.001, ****p<0.0001.

These results identify a major role for ESX-1 in driving an extensive neutrophilic inflammation at the expense of the monocyte-dominated low-grade inflammatory situation in ESX-1-deficient infection (Fig. 1H and S1A-B Fig.).

### Neutrophils and Ly6C^+^MHCII^+^ monocytes are the main cellular reservoirs for *M. marinum* in the infected tissue

To enable flow cytometry-based identification of infected cells in the tissue, WT and ΔRD1 bacteria were transformed with a plasmid encoding the green fluorescent protein ‘Wasabi’ (S2A Fig.). Although hygromycin – providing selective pressure to maintain the plasmid – was not administered *in vivo*, analysis of plasmid curing in infected mice demonstrated that it remained stable for at least 21 days post infection (S2B Fig.). During this time frame, the number of Wasabi-positive cells correlated with colony forming units (S2C Fig.). At 28 days post infection, plasmid curing occurred primarily in WT infection (S2B Fig.), which might reflect a higher rate of replication for WT bacteria compared to ΔRD1.

Bacteria were present in the CD45^+^ compartment (Fig. 2A-B and S2A Fig.), indicating that *M. marinum* resides in hematopoietic cells. While less than 5% of CD45^+^ cells were infected at any time point, these accounted for ∼100% of all infected host cells (Fig. 2C). Among CD45^+^ cells, neutrophils and CD64^+^ cells constituted the main Wasabi-positive populations (Fig. 2D-F) suggesting a limited role, if any, for additional cellular niches. Kinetic analysis suggested an increase in the number of infected neutrophils in WT infected animals over time (Fig. 2E), which may be explained by the concurrent ESX-1-mediated buildup of neutrophils in the tissue (Fig. 1C). The fraction of neutrophils that harbored bacteria was similar in WT and ΔRD1 infections (Fig. 2E), suggesting that ESX-1 did not affect the infectivity rate of recruited cells. The number of infected CD64^+^ cells (Fig. 2E) also largely correlated with their influx into the tissue (Fig. 1C, E), and ESX-1 had only a minor effect on the relative infection rate of these cells (Fig. 2E). Analysis of the CD64^+^ compartment suggested a role for G1 monocytes as a reservoir during the initial phase of infection, but that G2 monocytes rapidly became the dominating bacteria-harboring subset (Fig. 2G-H). Thus, neutrophils and Ly6C^+^MHCII^+^ inflammatory monocytes constitute the main cellular reservoirs for *M. marinum in vivo*.

**Fig. 2.**
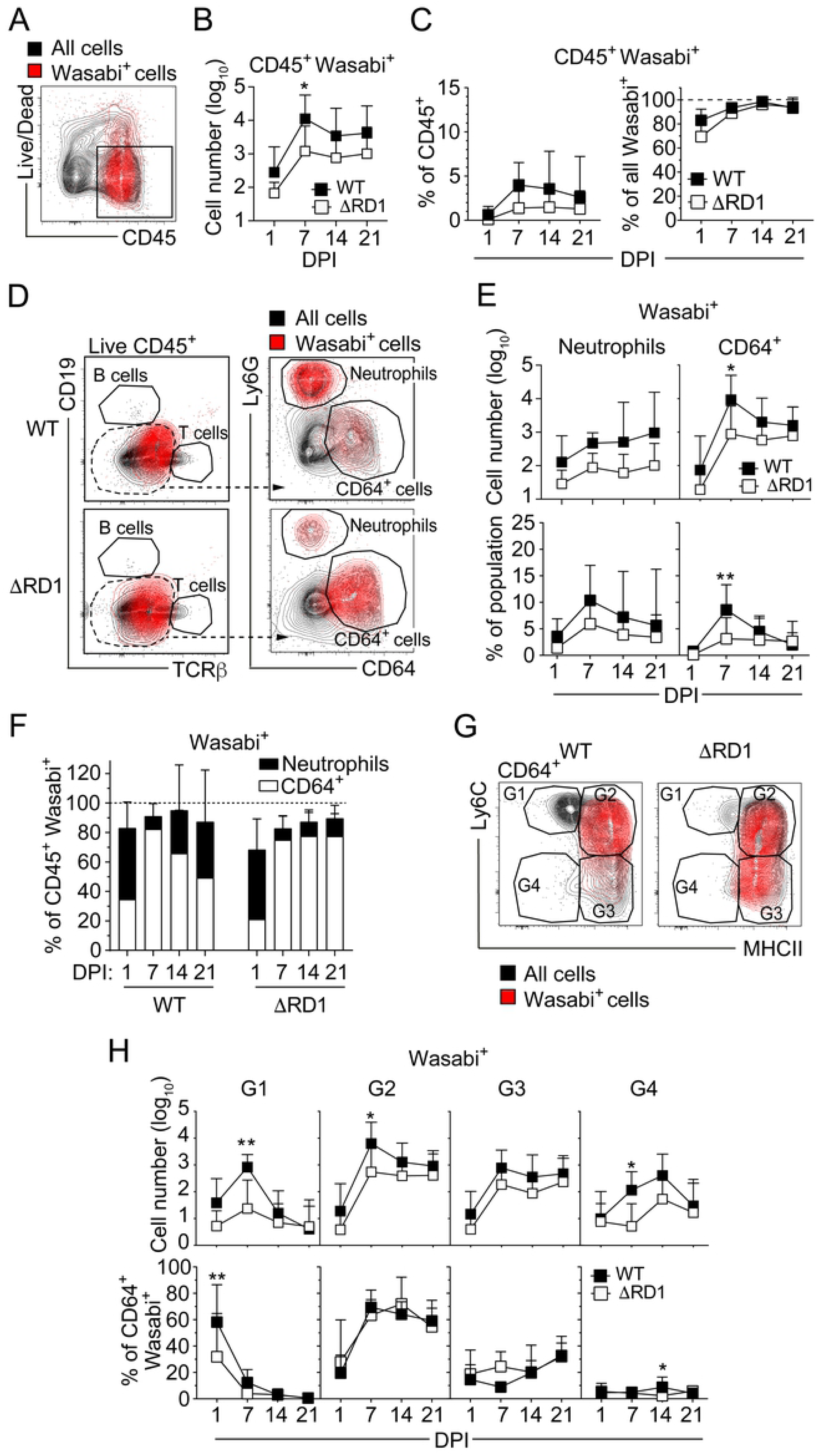
Neutrophils and Ly6C^+^MHCII^+^ monocytes are the main cellular reservoirs for *M. marinum* in the infected tissue. Flow cytometry analysis of tail tissues from C57Bl/6 mice infected with 5 x 10^6^ CFUs of Wasabi-expressing WT or ΔRD1 *M. marinum*. **A**) Representative flow cytometry plot of Wasabi expression among all cells in a WT infected mouse at 21 DPI. **B**) Quantification of live CD45^+^Wasabi^+^ cells at indicated DPI as gated in A. **C**) Proportion of Wasabi^+^CD45^+^ cells among total CD45^+^ cells (left panel) or among total Wasabi^+^ cells (right panel). **D**) Representative flow cytometry plots of Wasabi^+^ cells in the hematopoietic compartment. Live CD45^+^CD19^-^TCRβ^-^ cells were gated for analysis of Ly6G and CD64 expression. **E**) Numbers of Wasabi^+^ neutrophils and CD64^+^ as indicated (top panels) and their corresponding proportion among total neutrophils or CD64^+^ cells (bottom panels). **F**) Proportion of Wasabi^+^ neutrophils and CD64^+^ cells among total Wasabi^+^CD45^+^ cells. **G**) Representative flow cytometry plot of Wasabi^+^ cells among the G1 to G4 subpopulations of CD64^+^ cells. **H**) Numbers of Wasabi^+^ G1 to G4 populations (top panel) and their corresponding proportion among total Wasabi^+^CD64^+^ cells (bottom panel). (**A**-**H**) Results (mean + SD, *n* = 7-9 mice) from two independent experiments. 2-way ANOVA, *p<0.05, **p<0.01.

### ESX-1 promotes localized neutrophil accumulation in the granuloma core region

To examine the role for ESX-1 in the development of spatially organized granulomas we analyzed tail tissue in WT and ΔRD1 infected animals by multicolor fluorescence microscopy. Tail sections were collected at different time points post infection and stained with antibodies against Gr1 and CD64 to visualize neutrophils and monocytes/macrophages, respectively (Fig. 3). Bacteria were identified by their expression of Wasabi, and myeloperoxidase (MPO) was stained for as a marker of neutrophil activation (Fig. 3). The tail bone and bone marrow were identified by light microscopy, and as previously described [7] we observed ESX-1-dependent bone deterioration primarily after two weeks of infection (Fig. 3), reflecting the strong inflammatory milieu in WT infected tissues [7].

**Fig. 3.**
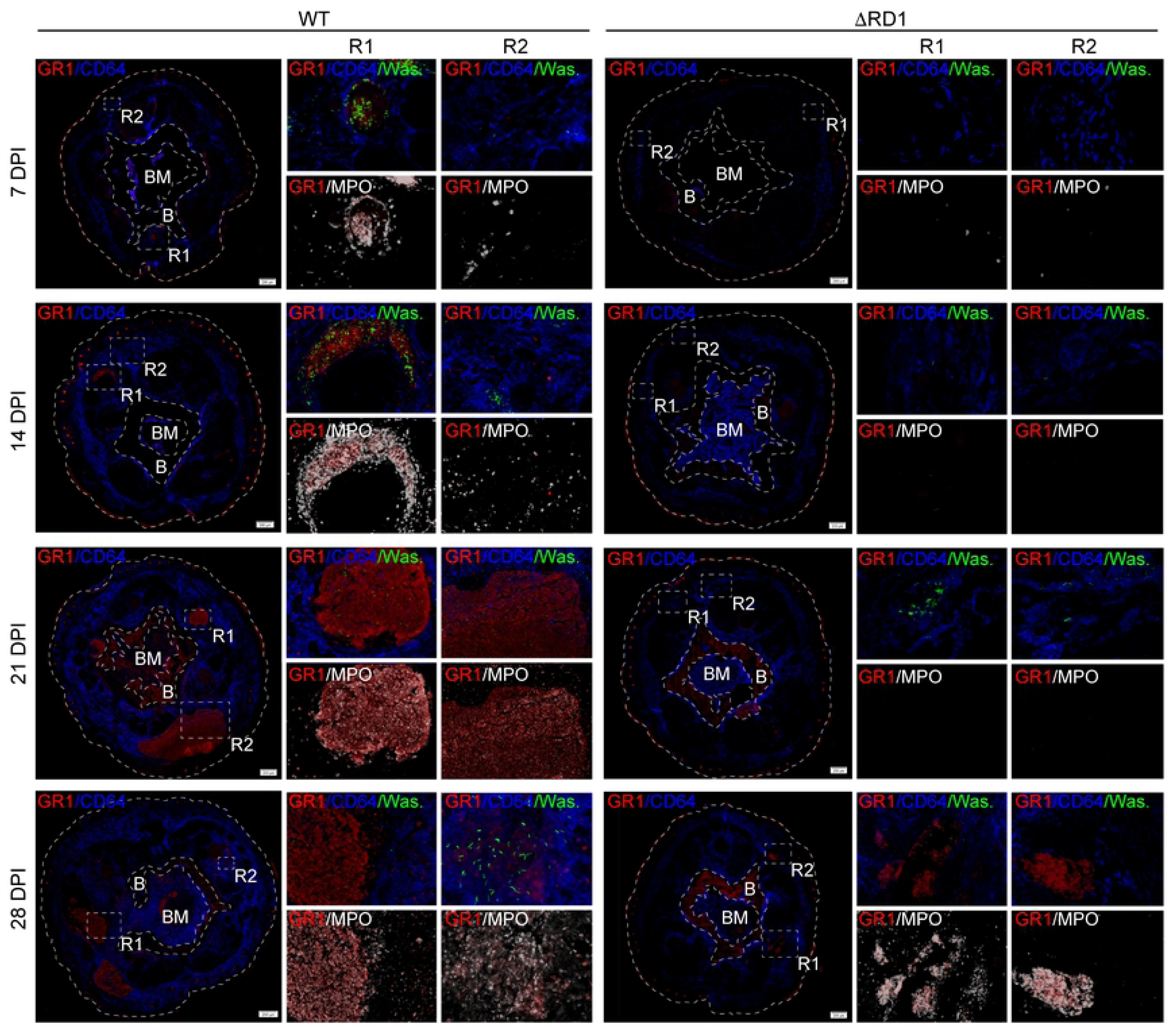
ESX-1 promotes localized neutrophil accumulation in the granuloma core region. Microscopy analysis of tail cross sections from C57Bl/6 mice infected with 5 x 10^7^ CFUs of Wasabi-expressing WT or ΔRD1 *M. marinum*, at the indicated DPI. Immunofluorescence staining of Gr1^+^ neutrophils (red), CD64^+^ cells (blue), MPO^+^ cells (white), and Wasabi^+^ bacteria (green). Boxes, R1 and R2, indicate enlarged regions shown in the right panels. Scale bar; 200 μm in left panel. Images were acquired using Z-stack acquisition with extended focus imaging (EFI) on an OLYMPUS VS-120 virtual slide microscope. R1, Region 1; R2, Region 2; B, Bone; BM, Bone marrow. Analysis performed on 2-3 tail cross sections per mouse (*n* = 3 mice per group per time point).

At 7 days post infection we observed ESX-1-dependent formation of small stratified granulomatous lesions, where bacteria localized primarily to central regions of MPO^+^ neutrophils that were surrounded by CD64^+^ cells (Fig. 3). The size of these nascent granulomas in WT infected animals increased over time and at all time points bacteria were observed mainly within the expanding neutrophilic core (Fig. 3), implying bacterial growth in this region. In contrast, ΔRD1 infection did not induce the formation of neutrophil-containing granulomatous structures until 28 days post infection, and ESX-1-deficient bacteria were mainly scattered in areas of CD64^+^ cells (Fig. 3). Of note, because WT bacteria showed significant curing of the Wasabi-encoding plasmid at 28 days post infection (fig. S2B), our microscopy-based analysis likely underestimates WT bacterial load at this time point (Fig. 3). Thus, ESX-1 promotes intra-granuloma accumulation of neutrophils, where they might serve to support bacterial growth.

### Neutrophils are required for ESX-1-mediated virulence

The role of neutrophils in mycobacterial infection is controversial [18-20, 22, 24-28]. In addition, results from previous antibody-based neutrophil depletion experiments are confounded by the recent discovery that injection of anti-Ly6G antibodies alone causes incomplete depletion [36], and that the extensively used anti-Gr1 antibody does not specifically target neutrophils [37]. To evaluate the functional role of neutrophils we therefore depleted these cells in WT infected mice by consecutive injections of rat anti-mouse Ly6G and mouse anti-rat κ light chain monoclonal antibodies (Fig. 4A) [36]. On average this treatment diminished the number of neutrophils by >90% throughout the infection (Fig. 4B). Treatment did not affect the number of CD64^+^ cells (Fig. 4B), but due to the loss of neutrophil accumulation they increased as a percentage of CD45^+^ cells by two weeks post infection (Fig. 4B).

**Fig. 4.**
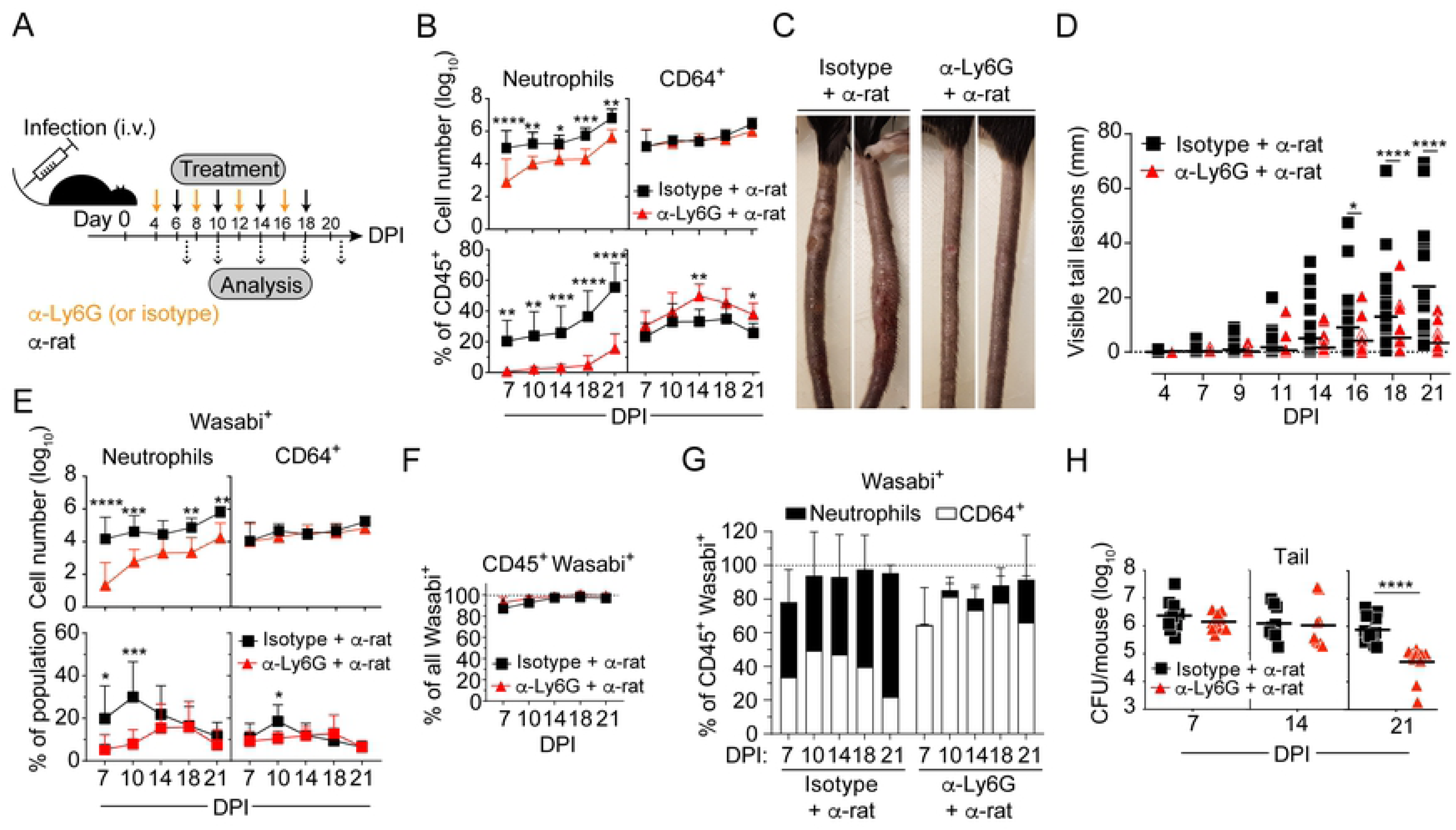
Neutrophils are required for ESX-1-mediated virulence. Neutrophil depletion in C57Bl/6 mice infected with 5 x 10^7^ CFUs of Wasabi-expressing WT *M. marinum*. **A**) Experimental set up for neutrophil depletion. Consecutive intraperitoneal injections of rat anti-mouse Ly6G (or isotype control) and mouse anti-rat *kappa* light chain monoclonal antibodies, as indicated. **B**, **E**-**G**) Flow cytometry analysis of tail tissues. **B**) Neutrophil and CD64^+^ cell numbers (top panels), and their corresponding proportions among total CD45^+^ cells (bottom panels). **C**) Visible tail lesions of representative mice at 21 DPI. **D**) Cumulative length of visible tail lesions. **E**) Numbers of Wasabi^+^ neutrophils and CD64^+^ cells (top panels) and their corresponding proportion among total neutrophils and CD64^+^ cells (bottom panels). **F**) Proportion of Wasabi^+^CD45^+^ cells among total Wasabi^+^ cells. **G**) Proportion of Wasabi^+^ neutrophils and CD64^+^ cells among total Wasabi^+^CD45^+^ cells. **H**) Bacterial burden in the tail. (**B**, **E**-**G**) Results (mean + SD, *n* = 9 mice) from three independent experiments. (**D**) Results (*n* ≥ 21 mice) from three independent experiments. Bars indicate the mean for each group. (**H**) Results (*n* = 9-12 mice; two-tailed unpaired t test) from two independent experiments. Bars indicate the mean for each group. (**B, D**-**F**) 2-way ANOVA. *p<0.05, **p<0.01, ***p<0.001, ****p<0.0001.

Analysis of visible lesions demonstrated drastically reduced disease-progression and inflammation in neutrophil-depleted animals (Fig. 4C-D), indicating that the extensive inflammation observed in ESX-1-proficient infection requires neutrophils.

Treatment reduced the number of bacteria-harboring neutrophils without causing a reciprocal increase of infected CD64^+^ cells (Fig. 4E). Cellular infection remained restricted to the CD45^+^ compartment in both neutrophil-depleted and control animals (Fig. 4F), and neutrophils and CD64^+^ cells accounted for essentially all infected cells in both experimental groups (Fig. 4G). Depletion of neutrophils translated into a reduced bacterial load only after two weeks of infection (Fig. 4H), trailing the kinetics of when these cells normally reach greater numbers (Fig. 1C and Fig. 3). Thus, following their accumulation in the tissue neutrophils support bacterial growth, which might be explained by their ability to generate a tissue environment permissive for extracellular growth [19]. The finding that bacterial load was unaffected for the first two weeks of infection in treated animals (Fig. 4H) implies that neutrophils themselves might not represent a major niche for intracellular replication. In agreement with this interpretation, *in vitro* studies suggested that *M. marinum* is less able to propagate in neutrophils of both mouse and human origin, compared to the intracellular growth observed in macrophages (S3 Fig.).

### ESX-1 drives differentiation of newly recruited uninfected neutrophils into a proinflammatory phenotype

To further explore how neutrophils mediate inflammation and to investigate if ESX-1, in addition to promoting neutrophil accumulation (Fig. 1 and Fig. 3), also regulates neutrophil phenotype, we performed single cell RNA-seq on infected and bystander (*i.e.* uninfected) neutrophils isolated from WT and ΔRD1 infected tail tissues two weeks post infection (Fig. 5A). After quality control and removal of contaminating cells, cellular debris and doublets from our sequencing data, we obtained single cell data for ≥3,242 neutrophils in each analytical group, with an average coverage of 866 genes per cell.

**Fig. 5.**
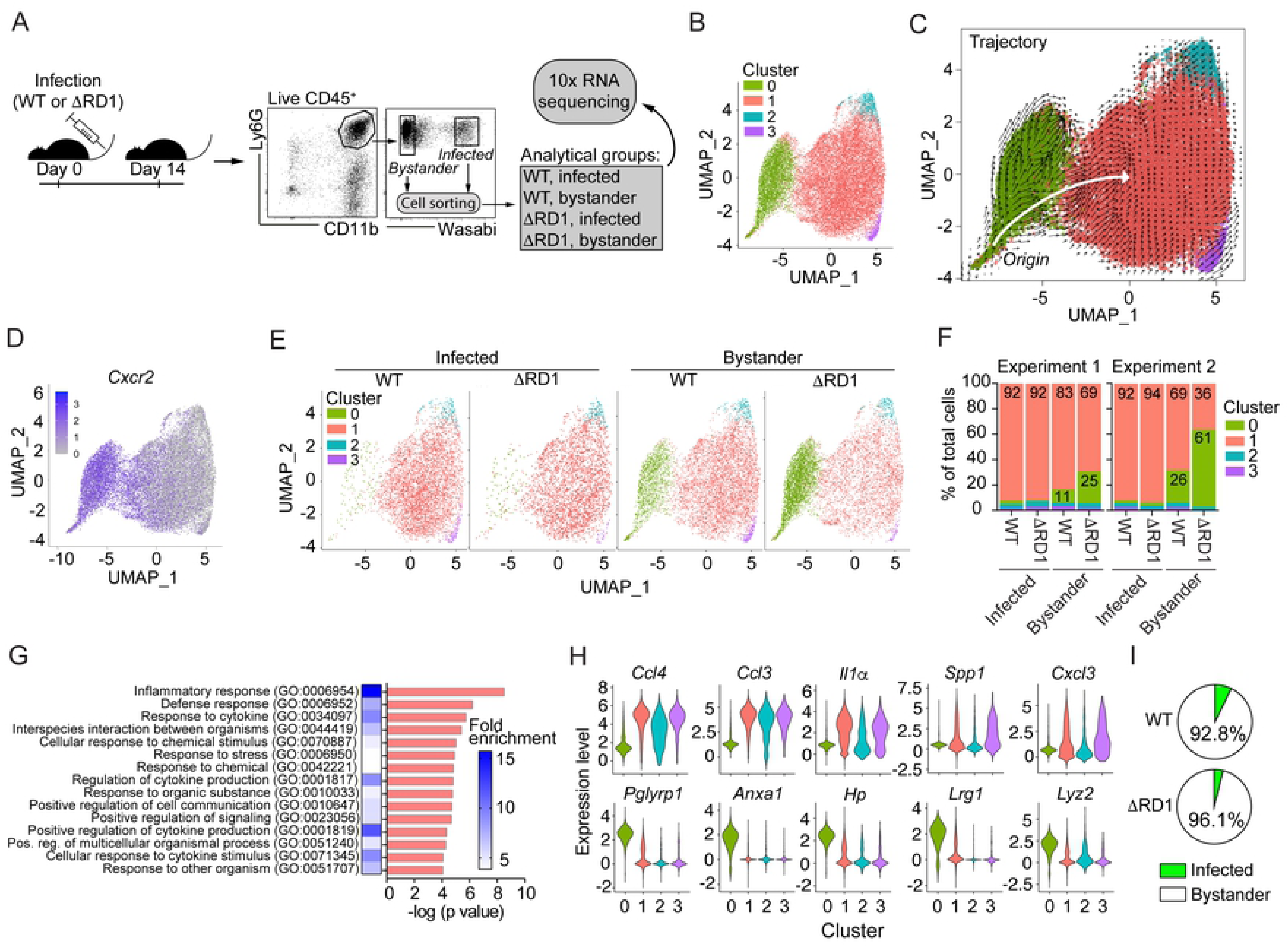
ESX-1 drives differentiation of newly recruited uninfected neutrophils into a proinflammatory phenotype. Single cell RNA-sequencing analysis of neutrophils sorted from tail tissues 14 DPI from C57Bl/6 mice infected with 5 x 10^7^ CFUs of Wasabi-expressing WT or ΔRD1 *M. marinum*. Results based on two independent experiments. **A**) Experimental set up of the single cell RNA-seq analysis using the 10X genomics approach. Infected (Wasabi^+^) and non-infected (Wasabi^-^) neutrophils (Live CD45^+^Ly6G^+^CD11b^+^) were sorted (15,000 cells per analytical group) by FACS and single-cell RNA-seq was performed. **B**) Louvain clusters at resolution 0.1 visualized on UMAP embedding. **C**) RNA velocity estimates for neutrophils shown on UMAP embedding of full data set and coloured by clustering from B. White arrow visualizes the overall direction of differentiation from origin. **D**) UMAP embedding with *Cxcr2* expression overlay. **E**) UMAP plot from B split into individual analytical groups as indicated. **F**) Cluster distribution for each analytical group in each of the two independent experiments performed. **G**) Gene ontology (GO) analysis (Biological processes) of cluster 1 compared to all other clusters and ordered by the -log (p value). The first 15 GO terms are indicated as well as their corresponding fold-enrichment score on the heat-map. **H**) Violin plots of single cell RNA expression level within clusters. Top 5 differentially expressed genes (DEGs) between cluster 1 (top row) or 0 (bottom row) and all other clusters, ordered by log fold-change values. **I**) Proportion of infected and uninfected bystander neutrophils in the infected tissue, as indicated. Results (mean, *n* = 7-9 mice) from two independent experiments.

Nearest neighbor analysis, using Louvain clustering at resolution 0.1 on pooled data from all analytical groups, identified four phenotypically distinct clusters (Fig. 5B), where clusters 0 and 1 accounted for ∼95% of all cells. To understand the relationship between the different phenotypes we performed velocyto-based trajectory analysis estimating RNA velocities. The main trajectory originated in cluster 0 from where cells differentiated into cluster 1 and the minor clusters 2 and 3 (Fig. 5C), suggesting that cluster 0 represents newly recruited non-differentiated neutrophils. Consistent with this interpretation CXCR2, which is required for neutrophil migration into infected tissues [20], was expressed in cluster 0 and downregulated upon differentiation (Fig. 5D).

Analysis of the individual analytical groups revealed that virtually all infected cells (≥92%) belonged to cluster 1 (Fig. 5E-F), implying that infected neutrophils are activated in an ESX-1-independent manner. In contrast, a considerable proportion of bystander cells had a non-differentiated phenotype, a feature that was accompanied by a corresponding decrease of cells in cluster 1 (Fig. 5E-F). ESX-1 reduced the fraction of non-differentiated cells >2-fold, suggesting that ESX-1 drives activation of newly recruited uninfected neutrophils into cluster 1 via an extrinsic mechanism (Fig. 5E-F). Gene ontology (GO) analysis based on ≥0.5-log fold differentially expressed genes (DEGs) indicated that cluster 1 cells have a strongly upregulated inflammatory phenotype as compared to cells in cluster 0 (Fig. 5G and S1 Table). Indeed, the most highly upregulated DEGs defining cluster 1 were proinflammatory cytokines and chemokines (Fig. 5H, top row and S1 Table). The expression of inflammatory mediators was paralleled by that of PDL1 and a downregulation of CD177 (S4A Fig.), implying phenotypic similarities between cluster 1 and protumorigenic neutrophils (reviewed in ref. [38, 39]). The minor clusters 2 and 3 separated from cluster 1 largely based on expression of ribosomal and heat shock proteins, respectively (S1 Table). Because the bystander population represents ≥90% of all neutrophils in the infected tissue (Fig. 5I), ESX-1-mediated differentiation of these cells into a proinflammatory phenotype likely impacts on inflammation and disease development.

Examination of the top 5 DEGs defining cluster 0 (Fig. 5H, bottom row) showed that antibacterial lysozyme (*Lyz2*) and peptidoglycan recognition protein 1 (*Pglyrp1*) – which has been linked to susceptibility to mycobacterial infection in cattle [40] – were downregulated upon differentiation. Moreover, differentiation caused downregulation of haptoglobin (*Hp;* Fig. 5H, bottom row) and a reciprocal upregulation of heme oxygenase-1 (*Hmox1*; S4B Fig.), which may increase the availability of free iron [41]. Thus, ESX-1-mediated differentiation of bystander neutrophils might also contribute to a more permissive environment for the bacteria.

### Monocytes suppress neutrophil accumulation and inflammation

To determine the role of monocytes in our model we took advantage of CCR2^-/-^ mice, which exhibit defective monocyte recruitment to inflamed tissues [33, 42]. *M. marinum* caused similar disease in CCR2^+/+^ and heterozygous CCR2^+/-^ mice (S5A Fig.), in which monocytes traffic normally [42], allowing us to use CCR2^+/-^ littermate controls for these experiments. Analysis of tail tissue from WT infected animals 14 days post infection confirmed that CCR2-deficiency significantly reduced the number of CD64^+^ cells (Fig. 6A), which decreased from ∼35% to <5% of CD45^+^ cells in CCR2^-/-^ (Fig. 6A). The diminished CD64^+^ compartment was due to reduced numbers of G1-G3 monocytes (Fig. 6B), while the number of G4 tissue macrophages remained similar between CCR2^-/-^ and CCR2^+/-^ littermates (Fig. 6B). CCR2-deficiency caused a significant increase in neutrophil numbers (Fig. 6A), which accounted for >80% of the CD45^+^ compartment in tail tissue of CCR2^-/-^ mice (Fig. 6A), indicating that monocytes normally inhibit neutrophil accumulation. Consistent with this, CXCL1, 2 and 5 levels were elevated in the infected tissue of CCR2^-/-^ compared to CCR2^+/-^ control (S5B Fig.).

**Fig. 6.**
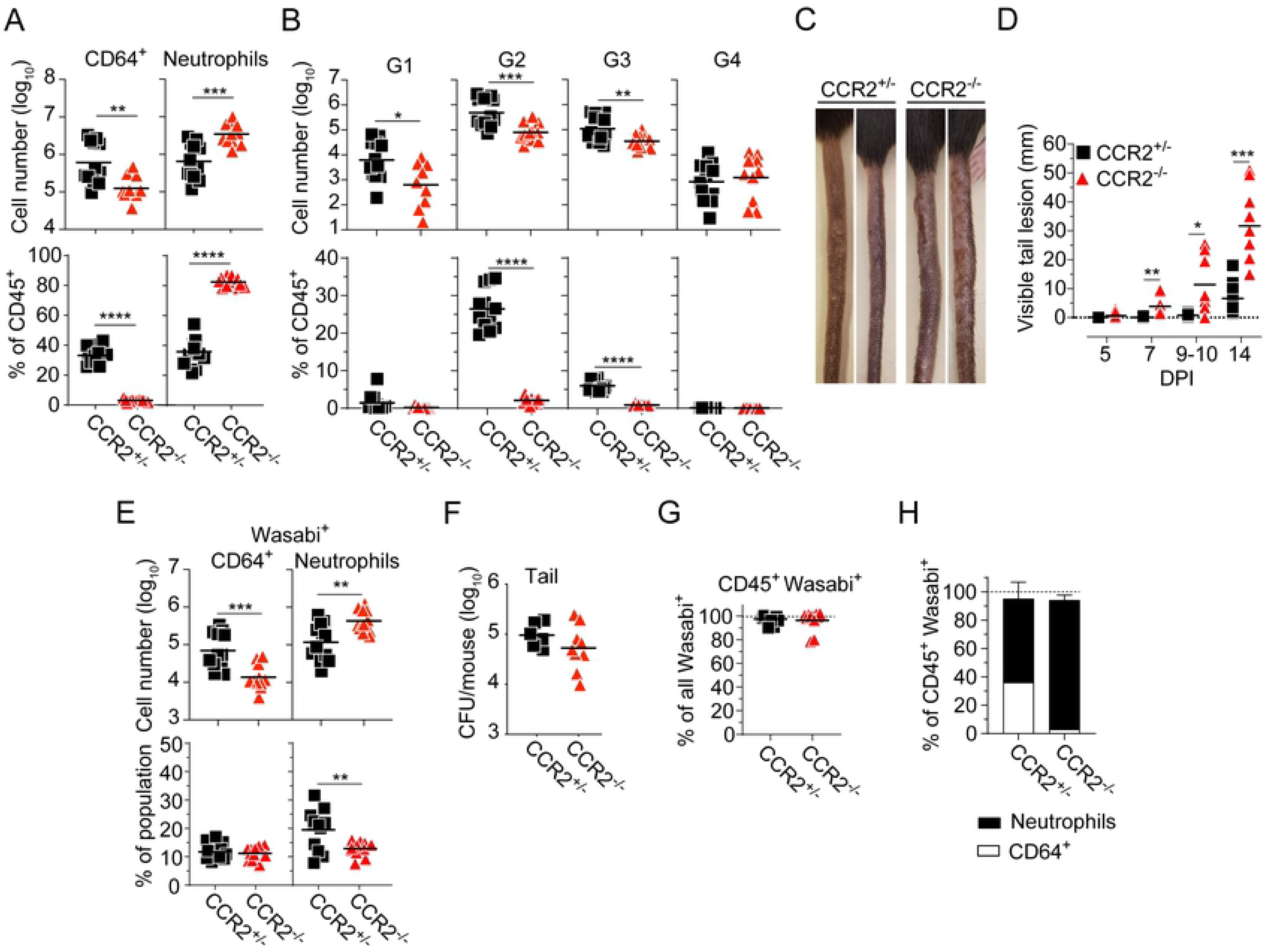
Monocytes suppress neutrophil accumulation and inflammation. CCR2^+/-^ or CCR2^-/-^ mice were infected with 5 x 10^7^ CFUs of Wasabi-expressing WT *M. marinum*. Unless indicated, analyses were performed at 14 DPI. **A**, **B**, **F**-**H**) Flow cytometry analyses of tail tissues. **A**) Numbers of CD64^+^ cells and neutrophils (top panels), as indicated, and their corresponding proportions among total CD45^+^ cells (bottom panels). **B**) Numbers of G1 to G4 CD64^+^ cells (top panels), and their corresponding proportions among total CD45^+^ cells (bottom panels). **C**) Visible tail lesions of representative infected mice. **D**) Cumulative length of visible tail lesions. **E**) Numbers of Wasabi^+^ neutrophils and CD64^+^ cells (top panels), and their corresponding proportion among total neutrophils and CD64^+^ cells (bottom panels). **F**) Bacterial burden in the tail. **G**) Proportion of Wasabi^+^CD45^+^ cells among total Wasabi^+^ cells. **H**) Proportions of Wasabi^+^ neutrophils and CD64^+^ cells among total Wasabi^+^CD45^+^ cells. (**A**, **B**, **F**, **G**) Results (*n* = 11-12 mice; two-tailed unpaired t or Mann-Whitney test) from three independent experiments. (**D**) Results (*n* = 11-12 mice; 2-way ANOVA) from three independent experiments. (**E**) Results (*n* = 6-8 mice; two-tailed unpaired t test) from two independent experiments. (**H**) Results (mean + SD, *n* = 11-12 mice per group) from three independent experiments. (**A**, **B**, **D**-**G**) Bars indicate the mean for each group. *p<0.05, **p<0.01, ***p<0.001, ****p<0.0001.

Infected CCR2^-/-^ mice exhibited dramatically increased inflammation and disease-progression (Fig. 6C-D). In fact, the development of highly inflamed and purulent tail lesions was so severe that CCR2-deficient mice often had to be sacrificed at two weeks post infection for ethical reasons, precluding analysis beyond this time point. These results (Fig. 6A-D) suggest a major host-protective function for monocytes by actively restricting ESX-1- mediated neutrophilic immunopathology.

The loss of monocytes as a cellular reservoir in CCR2^-/-^ mice (Fig. 6E) did not lead to significantly reduced bacterial load in the tail (Fig. 6F), which might be explained by the elevated numbers of neutrophils (Fig. 6A, E) that can promote a growth-permissive tissue environment [19]. Thus, monocytes limit disease (Fig. 6C-D) without affecting bacterial load (Fig. 6F), implying that they support disease tolerance to infection. Cellular infection was confined to the CD45^+^ compartment in both experimental groups (Fig. 6G), where neutrophils accounted for ∼95% of all infected cells in CCR2^-/-^ mice (Fig. 6H). Of note, the reduced percentage of infected neutrophils in CCR2^-/-^ (Fig. 6E) is likely due to the significantly increased total number of neutrophils in the tissue of these animals (Fig. 6A).

### iNOS activity in monocytes is required for their ability to suppress neutrophilic inflammation

Previous studies indicate a role for iNOS-derived nitric oxide in suppressing neutrophil recruitment to the site of *M. tuberculosis* infection [19, 21], making it of interest to determine if monocyte-mediated inhibition of neutrophil accumulation occurs via an iNOS-dependent mechanism. To test this hypothesis we first sought to define the cellular source of iNOS-derived nitric oxide in the infected tissue. Cell suspensions were prepared from WT and ΔRD1 infected tail tissues, and iNOS expression was determined by intracellular staining. *M. marinum* selectively induced iNOS in CD64^+^ myeloid cells essentially in an ESX-1-independent manner (Fig. 7A-B). While a higher proportion of the infected CD64^+^ cells expressed iNOS (Fig. 7C), uninfected cells constituted the largest iNOS expressing population by virtue of their abundance (Fig. 7C-D). Subdivision of the CD64^+^ population demonstrated that G2 monocytes accounted for ∼80% of all iNOS expressing cells in both WT and ΔRD1 infection (Fig. 7E-F), indicating that Ly6C^+^MHCII^+^ monocytes are the main producers of iNOS-derived nitric oxide in the infected tissue. In agreement with these results, infection of heterozygous CCR2-RFP mice, in which one of the CCR2 alleles is exchanged to express red fluorescent protein (RFP), indicated that >95% of all iNOS staining was within the CCR2^+^ compartment (Fig. 7G-H). Transcription of *Nos2* (iNOS) was significantly reduced in CCR2^-/-^ mice (Fig. 7I), confirming that monocytes are the major source of iNOS in the infected tissue.

**Fig. 7.**
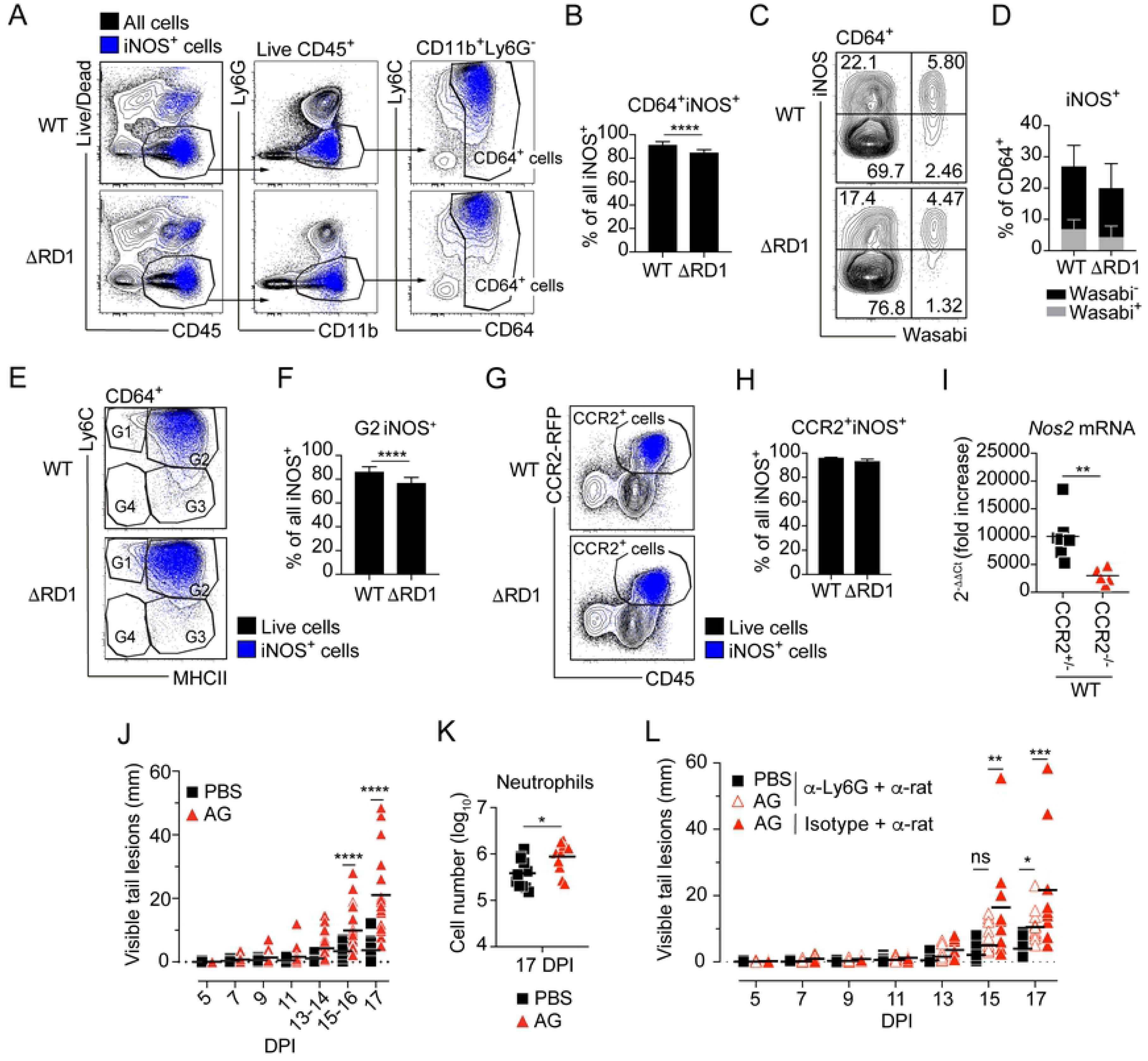
iNOS activity in monocytes is required for their ability to suppress neutrophilic inflammation. C57Bl/6 wild type (**A**-**F**, **J**-**L**), CCR2-RFP (**G**, **H**), and CCR2^+/-^ or CCR2^-/-^ (**I**) mice were infected with 5 x 10^7^ CFUs of Wasabi-expressing WT or ΔRD1 *M. marinum*, as indicated. **A**-**H**) Flow cytometry analysis of tail tissues at 14 DPI. **A**) Representative FACS plots, and **B**) quantification of iNOS^+^CD64^+^ cells. **C**) Representative FACS plots of iNOS expression in infected (Wasabi^+^) and uninfected (Wasabi^-^) CD64^+^ cells and **D**) the corresponding quantified data. **E**) Representative FACS plots of iNOS^+^ expression in the G1 to G4 gates of CD64^+^ cells. **F**) Proportion of iNOS^+^ G2 cells among total iNOS^+^ cells. **G**) Representative FACS plots of CCR2-RFP^+^ and CD45^+^ expression among iNOS^+^ cells and **H**) the corresponding quantified data. **I**) RTqPCR-based analysis of *Nos2* mRNA expression in tail tissues of WT infected CCR2^+/-^ or CCR2^-/-^ mice at 14 DPI. **J**-**L**) Treatment with aminoguanidine (AG) in WT infected C57Bl/6 mice. **J**) Cumulative length of visible tail lesions. **K**) Neutrophils numbers in tail tissues. **L**) Cumulative length of visible tail lesions in AG treated mice with or without neutrophil depletion as indicated. (**A**-**F**) Results (mean + SD, *n* = 12-13 mice; [**B**, **F**] two-tailed Mann-Whitney test, [**D**] 2-way ANOVA) from two independent experiments. (**G**, **H**) Results (mean + SD; *n* = 3 mice per group; two-tailed Mann-Whitney test). (**I**) Results (*n* = 5-8 mice; two-tailed Mann-Whitney test). (**J**) Results (*n* = 22-23 mice; 2-way ANOVA) from two independent experiments. (**K**) Results (*n* = 11 mice; two-tailed unpaired t test) from two independent experiments. (**L**) Results (*n* = 9-12 mice; 2-way ANOVA). (**I**-**L**) Bars indicate the mean for each group. *p<0.05, **p<0.01, ***p<0.001, ****p<0.0001; ns, not significant.

To determine if iNOS is required for the ability of monocytes to suppress neutrophilic inflammation we administered aminoguanidine (AG), which selectively inhibits iNOS activity [19], to WT infected mice. Similar to the situation in CCR2-deficient mice (Fig. 6 and S5B Fig.), pharmacological inhibition of iNOS enhanced both disease-development (Fig. 7J) and neutrophil accumulation (Fig. 7K), which correlated with elevated levels of CXCL1, 2 and 5 in the infected tissue (S5C Fig.). Concomitant depletion of neutrophils largely reversed the excessive disease observed in AG treated animals (Fig. 7L), demonstrating that the phenotype was caused by increased neutrophil accumulation. The modest effect of AG on disease in anti-Ly6G antibody-treated animals (Fig. 7L) might be due to reduced efficiency of neutrophil depletion in AG treated mice (S5D Fig.). These findings identify iNOS-mediated suppression of ESX-1-mediated neutrophil accumulation as a key mechanism by which monocytes exert their anti-inflammatory function in mycobacterial infection.

## Discussion

Recent studies of *M. tuberculosis* infected mice have identified neutrophils and monocyte-derived cells as the main cellular reservoirs in lung tissue [15–18], but the fates of bacteria in these different cell types remain elusive. While neutrophils have been reported to represent a niche for intracellular replication in tuberculosis [14, 18], direct evidence for significant intracellular replication in neutrophils is lacking. Monocyte-derived interstitial macrophages are relatively restrictive for *M. tuberculosis* growth due to a low cellular availability of ferrous iron as well as to a metabolic bias towards glycolysis over fatty acid oxidation [16]. Mycobacteria can also grow extracellularly in the tissue, a feature linked to dysregulated inflammatory responses in *M. marinum* infected zebrafish [43, 44]. We identify neutrophils and Ly6C^+^MHCII^+^ monocytes as the main bacteria-harboring cell types in *M. marinum* infected tissues. Neutrophils supported bacterial growth following their accumulation, but our results imply a minor role for neutrophils as a niche for intracellular replication. This is in agreement with the notion that a more inflammatory milieu – as generated by ESX-1- mediated neutrophil accumulation in our model – is associated with extracellular growth [43], and with the finding that neutrophils produce an iron- and nutrient-replete tissue environment permissive for *M. tuberculosis* replication [19].

Work with evolutionarily distant infectious agents, including *M. tuberculosis* [19, 21], *Streptococcus pneumoniae* [45], *Streptococcus pyogenes* [46, 47], and influenza [48] suggest a role for IL1 in neutrophil recruitment. As for neutrophils, IL1 has been ascribed both host-protective [49] and host-detrimental [19] roles in *M. tuberculosis* infection, an apparent paradox that might relate to disparate roles during different phases of infection or to the use of different experimental conditions. Neutrophil recruitment to *M. tuberculosis* infected tissues requires NLRP3-dependent IL1-signaling [19–21], and we have shown that *M. marinum* drives host-detrimental inflammation by ESX-1-dependent activation of the NLRP3/ASC inflammasome [7]. Here we find that ESX-1 promotes production of CXCL1, 2 and 5 and the accumulation of neutrophils in granuloma core regions. Strikingly, we demonstrate that ESX-1-mediated pathology is dependent on neutrophils, indicating that neutrophils are host-detrimental in mycobacterial infection and that they have a previously unrecognized required role in executing ESX-1-mediated virulence. We speculate that ESX-1-mediated and neutrophil-dependent immunopathology may promote caveation of granulomas and thus bacterial transmission to new hosts.

In addition to their known antibacterial functions, neutrophils can regulate inflammatory responses [50] and are emerging as key cells also in autoimmune diseases and cancer. The phenotype of bacteria-harboring and bystander neutrophils from the site of an infection has not previously been analyzed at a single cell level. However, neutrophil phenotype has been extensively studied in the context of tumors [38, 39], which have immunological similarities to mycobacterial granulomas. Similar to tuberculosis, high infiltration of neutrophils is associated with poor prognosis in many human tumors [39], and, analogously to our results, antibody-mediated depletion of neutrophils can protect from primary tumorigenesis and metastasis [51, 52]. ESX-1 promoted accumulation of neutrophils in the granuloma core region, akin to the intra-tumoral localization typically observed for protumorigenic neutrophils [39]. Neutrophils can, however, adopt both pro- and antitumorigenic properties [38, 39], and single cell RNA-seq analysis of tumor-associated neutrophils has revealed a spectrum of distinct phenotypes within the tumor tissue [38, 53]. Our results indicate that ESX-1 drives newly recruited uninfected bystanders into an inflammatory phenotype with parallels to neutrophils that promote tumor inflammation and growth, often characterized by expression of cc-chemokines and PDL1, and an ability to regulate both CD64^+^ myeloid cells and T cell responses [38, 39]. Thus, in addition to driving neutrophil accumulation, ESX-1 is required for an extrinsic mechanism by which bystanders – corresponding to over 90% of all neutrophils in the infected tissue – adopt a phenotype that might contribute to immunopathology and granuloma growth.

We show that monocytes protect against ESX-1-mediated host-detrimental neutrophilic inflammation without affecting bacterial load, demonstrating a direct immunosuppressive and disease tolerizing function of monocytes in mycobacterial infection. Unlike neutrophils, monocytes were recruited to the infected tissue in an ESX-1-independent manner; work in the zebrafish model similarly suggests ESX-1-dependent neutrophil recruitment [28] whereas monocytes exhibit ESX-1-independent recruitment that depends on phenolic glycolipids of the mycobacterial cell wall [54, 55]. It was recently shown that iNOS-derived nitric oxide can inhibit neutrophil recruitment to *M. tuberculosis* infected tissues [19, 21]. While the cellular source of iNOS remained unknown, nitric oxide was found to limit IL1-mediated neutrophil recruitment by inhibition of the NLRP3 inflammasome [21]. Our findings identify Ly6C^+^MHCII^+^ monocytes as the main iNOS-expressing cell type in the infected tissue and, importantly, we demonstrate that iNOS activity is required for the mechanism by which monocytes suppress ESX-1-mediated immunopathology.

Collectively these results put the regulation of neutrophils at center stage of ESX-1- mediated virulence. Moreover, they reveal a yin and yang-like interplay between monocytes and neutrophils that may regulate immunopathology not only in mycobacterial infection, but also in other infections as well as in sterile inflammatory conditions and cancer.

## Materials and Methods

### Ethical statement

All animal care and use were in accordance with the Swedish animal welfare laws and guidelines from the Swedish Department of Agriculture (Act 1988:534). This work was approved by the Malmö/Lund Ethical Board for Animal Research, Lund/Malmö, Sweden (5.8.18-04144/2018 and 5.8.18-08454/2020).

### Animals

Female wild type C57Bl/6JRj mice were purchased from Janvier Labs. CCR2-deficient (B6.129S4-*Ccr2^tm1Ifc^*/J), CCR2^RFP^ (B6.129(Cg)-*Ccr2^tm2.1Ifc^*/J) and C57Bl/6NCrl mice were bred and maintained at the Lund University Clinical Research Center (CRC), Malmö. B6.129S4-*Ccr2^tm1Ifc^*/J and B6.129(Cg)-*Ccr2^tm2.1Ifc^*/J were originally purchased from The Jackson Laboratory. B6.129S4-*Ccr2^tm1Ifc^*/J were crossed with C57Bl/6NCrl mice, originally obtained from Charles River lab, to generate CCR2^+/-^ mice for CCR2^+/-^ x CCR2^-/-^ breeding. Littermate controls were used for experiments involving CCR2^+/-^ and CCR2^-/-^ mice.

### Bacterial strains and growth conditions

The wild type *M. marinum* M strain and its isogenic mutant (ΔRD1) lacking the RD1 locus [9] were used. Both strains carry the pTEC15 plasmid (Addgene) for green fluorescence (Wasabi) expression (selection with 50 µg/mL Hygromycin B). These strains, along with all herein used reagents, are listed in table S2. Bacteria were grown at 30°C in Middlebrook 7H9 medium (BD Biosciences) supplemented with 0.5% glycerol (Sigma-Aldrich), 0.05% Tween 80 (Sigma-Aldrich) and 10% ADC Supplement (BD Biosciences), or on Middlebrook 7H10 agar (BD Biosciences) supplemented with 0.5% glycerol and 10% OADC (Conda Lab).

### Mouse infections

7H9 culture medium was inoculated from frozen stocks and bacteria were grown until late log phase. Bacteria were washed twice in sterile PBS and bacterial clumps were dissociated by 3 passages through a 26G needle, followed by centrifugations at 500 g for 1 min to collect single-cell bacterial suspensions. Female mice between 8-12 weeks of age were infected via tail vein injections with 5 x 10^6^ or 5 x 10^7^ bacteria per mouse, as indicated in figure legends, in a total volume of 200 µL. Disease development was followed and quantified by measuring the length of individual visible skin lesions on the tail of each mouse. The cumulative length of all lesions per mouse tail was calculated and presented in millimeters.

### Neutrophil depletion

Mice were injected intraperitoneally (ip) with 100 µg of either the rat *InVivo*MAb anti-mouse Ly6G monoclonal antibody (BioXell, clone 1A8) or the rat anti-trinitrophenol isotype control (BioXell, clone 2A3) followed by an injection 48 h after with 100 µg of a mouse anti-rat immunoglobulin κ light chain (BioXell, clone MAR18.5). This procedure was repeated 4 times. Neutrophil depletion using the anti-Ly6G antibody has been shown to induce membrane antigen masking on depletion escaping cells [36]; therefore, neutrophils were identified as CD11b^+^Gr1^high^CD64^-^ in these experiments.

### Aminoguanidine (AG) treatment

Mice were injected ip with 200 µL of 50 mg/mL AG hemisulfate salt (Sigma-Aldrich) resuspended in PBS from day 1 post infection and every 48 h for a total of 8 injections. When neutrophil depletion was performed in AG treated animals, neutrophil neutralizing antibodies (see neutrophil depletion section above) were mixed with the AG solution for ip injections at day 3, 5, 7, 9, 11, 13 and 15 post infection.

### Analysis of bacterial burden

Spleens, tails, as well as the sciatic, inguinal and mesenteric lymph nodes were collected. Spleen and lymph nodes were disrupted in PBS 0.1% Triton X100 using a stainless bead with the TissueLyser II (Qiagen). The tail-draining sciatic and inguinal lymph nodes were combined before disruption. Tails were severed from mice at the tail base and cut into 3 mm pieces, which were homogenized in 3 mL PBS 0.1% Triton X100 using a homogenizer PT 1200 E (Polytron). Homogenized tissues were serially diluted and plated on 7H10 agar plates for enumeration of colony forming units (CFUs).

### Flow cytometry analysis

Tails were severed from mice at the tail base, and the tissue was separated from the bones after a longitudinal excision. Samples were first cut into 2 mm pieces, transferred in DMEM supplemented with 5% fetal calf serum (FCS), 30 µg/mL Liberase TM (Roche) and 52 µg/mL DNAse I (Sigma), and subsequently incubated with magnetic stirring for 60 min at 37°C. Samples were mashed through a 70 µm nylon cell strainer, washed in FACS buffer (PBS supplemented with 3% FCS and 2 mM EDTA) and filtered through a 40 µm nylon cell strainer. To block Fc-receptors, cells were first incubated with a rat anti-mouse CD16/CD32 antibody (clone 2.4G2). Cell surface staining was performed in PBS for 30 min on ice with the fixable viable dye Near-IR Dead Cell Stain Kit (Invitrogen) using the following fluorochrome-conjugated anti-mouse antibodies: CD45.2 (clone 104 or 30F-11), CD11b (clone M1/70), CD11c (clone N418), CD64 (clone X54-5/7.1), Ly6C (clone HK1.4), Ly6G (clone 1A8), MHCII (clone M5/114.15.2), Gr-1 (clone RB6-8C5), CD19 (clone 1D3 or 6D5), TCRβ (clone H57-597), CD4 (clone GK1.5 or RM4-5), CD8a (clone 53-6.7), CD3 (clone 17A2), Nos2 (clone C-11) or its mouse IgG1k isotype control (clone MOPC-21). Subsequently, cells were fixed with 2% PFA for 20 min at RT. AccCount Fluorescent Particles (Spherotech) were added to each sample to evaluate the total number of cells in the sample during flow cytometry analysis. Flow cytometry analyses were performed on an LSR II flow cytometer (BD Biosciences), and data were analyzed using the FlowJo software versions 9 and 10. Intracellular iNOS staining was performed after cell surface marker staining and fixation as described above. Cells were permeabilized in permeabilization buffer from the FoxP3 staining buffer set (eBiosciences) in the presence of 2% rat serum for 15 min, and intracellular staining was performed overnight at 4°C in permeabilization buffer/rat serum.

### Immunofluorescence microscopy analysis

Tails were severed from the mice ∼1 cm from the tail base, and 2-3 conjunctive pieces of ∼0.5-1 cm long were collected at the severed end. Tail pieces were fixed in 3% AntigenFix (Diapath) for 2 h on ice, and then incubated in PBS supplemented with 35% sucrose on rotation over-night at 4℃. Dehydrated samples were frozen in Tissue-Tek OCT and cut into 15 μm cross sections with a CM1950 cryostat (Leica). Cryostat sections were incubated in permeabilization buffer (PBS containing 1% saponin, 2% BSA, 1% FCS, 1% donkey serum) supplemented with 1% mouse serum for 30 min at RT. For staining, tissues were incubated over night at 4℃ in permeabilization buffer with purified polyclonal rabbit anti-mouse myeloperoxidase (MPO) IgG (Thermo Fisher; PA5-16672) and AF647-conjugated mouse anti-mouse CD64 IgG1 (clone X54-5/7.1). MPO was visualized with secondary alpaca anti-rabbit AF-594-conjugated IgG (Jackson ImmunoResearch; 615- 585-214) for 1 h at RT. After blocking in permeabilization buffer supplemented with 2% rat serum for 30-60 min at RT, sections were stained with a Pacific Blue™-conjugated rat anti-mouse Gr1 antibody (clone RB6-8C5) for 1 h at RT. Tissues were subsequently washed and mounted in ProLong Gold (Invitrogen). Images were acquired using Z-stack acquisition with extended focus imaging (EFI), using a 20X objective on an OLYMPUS VS-120 virtual slide scanning microscope and the Olympus VS-ASW-S6 software, and processed using Photoshop 2020/2021 (Adobe).

### ELISA and RTqPCR analysis

Tails were severed from mice at the tail base. The tissue was separated from the bone after a longitudinal excision and was immediately stored at -80°C. Frozen tissue was immersed in liquid nitrogen and pulverized with a biopulverizer (Biospec Products) that had been pre-chilled in liquid nitrogen. Pulverized tissue samples were transferred to PBS containing Complete EDTA-free protease inhibitor cocktail (Roche), and homogenized with a homogenizer PT 1200 E (Polytron). Homogenized tissues were either processed for ELISA analysis or RNA extraction.

For ELISA analysis, samples were incubated 90 min on ice and then centrifuged at 500 g (10 min at 4°C). Supernatants were collected and further centrifuged at 20,000 g (15 min at 4°C) to pellet remaining debris. Finally, supernatants were collected and analyzed for mouse CXCL1/KC, mouse CXCL2/MIP-2 and mouse CXCL5/LIX using the corresponding ELISA kits (RnD Systems).

RNA was extracted using the RNeasy Mini Kit (Qiagen) with DNase digestion. RNA concentrations were quantified using a Nanodrop ND1000 (Thermo Fisher), and were normalized across samples. cDNA was generated using the GoScript Reverse Transcription System (Promega). RTqPCR analysis was conducted in 384-well format using the CFX 384 Real-Time system (BioRad) and SoFast EvaGreen qPCR super mix (BioRad). All infected samples were normalized to expression of the housekeeping gene *reep5* as well as to corresponding uninfected samples. The following primer sequences were used: *Nos2* forward (5′-TGGAGCGAGTTGTGGATTGTC) and *Nos2* reverse (5′-GGGCAGCCTCTTGTCTTT GA); *reep5* forward (5′-GCCATCGAGAGTCCCAACAA) and *reep5* reverse (5′-GCATCT CAGCCCCATTAGC).

### Single cell RNA-sequencing

Two independent experiments were performed. For each experiment 14 to 15 tails per group were collected, pooled and prepared as described for flow cytometry analysis. Live CD45^+^Ly6G^+^CD11b^+^Wasabi^+^ or Wasabi^-^ cells were sorted by FACS in PBS supplemented with 1% BSA using a FACS ARIA Fusion (BD Biosciences). Single cell RNA-sequencing libraries were prepared according to the manufacturer’s instructions using Chromium Single Cell 3′ Library & Gel Bead Kit v3 (10x Genomics, PN-1000092) and Chromium Chip B Single Cell Kit (PN-1000074) with the Chromium Controller & Next GEM Accessory Kit (10x Genomics, PN-120223). In brief, single cells, reverse transcription reagents, Gel Beads containing barcoded oligonucleotides, and oil were combined on a microfluidic chip to form Gel Beads in Emulsion (GEMs). Individual cells were lysed inside distinct GEMs and the released poly-A transcripts were being barcoded with an Illumina R1 sequence, a 10X barcode and a Unique Molecular Identifier (UMI) during the reverse transcription (RT). After RT, the GEMs were broken, the barcoded cDNA were purified using Dynabeads MyOne silane (10x Genomics, PN-2000048) and amplified by PCR. Amplified cDNAs were cleaned up with SPRIselect Reagent kit (Beckman Coulter, B23318). Indexed sequencing libraries were constructed by enzymatic fragmentation, end-repair and A-tailing, before a second and final PCR amplification using the Chromium i7 Sample Index (10x Genomics, PN-220103), introducing an Illumina R2 sequence, a unique sample index (allowing multiplex sequencing) and P5/P7 Illumina sequencing adaptors to each library. Library quality control and quantification were performed using a KAPA Library Quantification Kit for Illumina Platforms (Kapa Biosystems, KK4873) and the 2100 Bioanalyzer equipped with a High Sensitivity DNA kit (Agilent, 5067-4626).

Multiplexed libraries were pooled and Illumina deep sequencing was performed either by NextSeq 500/550 High Output v2.5 kit (150 cycles) at the Center of Excellence for Fluorescent Bioanalytics (KFB, University of Regensburg, Germany) or by Novaseq 6000 S1 or S2 (200 cycles) at the SNP&SEQ Technology Platform (Uppsala, Sweden) with the following sequencing run parameters: Read1 - 28 cycles; i7 index - 8 cycles; Read2 – 126 cycles at a depth of at least 100M reads/sample.

The single cell RNA-seq data are deposited in NCBÍs Gene Expression Omnibus (GEO) and are accessible through GEO series accession number GSE172072.

### Bioinformatic analyses of single cell RNA-seq data

After retrieval, the sequencing data was processed and aligned using CellRanger (version 2.1.1 for the first experiment and version 3.1.0 for the second independent experiment) [56, 57] using mm10 as reference for the first experiment and mm10-2.1.0 for the second. The data was then read into R (version 3.5.1 and 4.0.2) and each sample quality checked by mitochondrial gene content, number of reads and genes. Cells with very low counts of reads and genes, and with high mitochondrial content were discarded from further analysis. The experimental replicates for each analytical group were normalized, integrated, variable genes calculated (with vst), expression scaled, dimensionality reduced with PCA, and cells clustered with Louvain clustering all using Seurat[58]. Cell cycle associated gene expression per cell (genes from [59]) was calculated with AddModuleScore and regressed out during scaling of the gene expression with linear regression. Numbers of PCs for further analyses were decided in individual models by ElbowPlot and DimHeatmap. The individual analytical groups were then analyzed to exclude doublets (cells clustering due to high or low gene and read content) and contaminating cells such as erythrocytes which were recognized based on differentially expressed genes specific for the cell type (*Bpgm*, *Hba-a1*, *Hba-a2*, *Hbb-bt*, *Hbb-bt*, *Fech*, *Gypa*, *Alas2*) [60]. Differential gene expression was calculated with Seurats FindAllMarkers using non-parametric Wilcoxon rank sum test. The remaining cells from each analytical group were integrated into one data set which was dimensionality reduced first with PCA then with UMAP, and reclustered.

The initial part of velocyto analysis was run on the outputs from CellRanger. The output loom files from the WT bystander group were read into R and combined into two matrices, one with spliced and one with unspliced data. From the velocyto [61] library for R armaCor was used to calculate cell-to-cell distances based on PCA embedding and filter.genes.by.cluster.expression to remove genes expressed below 0.2 for spliced data and 0.05 for unspliced data. The relative velocities were calculated with deltaT=1, kCells=30 and fit.quantile=0.02. Finally, the relative velocities were projected onto the existing UMAP embedding of all four analytical groups with n=200, scale=sqrt, arrow.scale=2, min.grid.cell.mass=0.5 and grid.n=40.

Gene ontology (GO) analysis was performed with the Panther Overrepresentation test (version DOI: 10.5281/zenodo.4033054) using the GO Biological process annotation data set [62]. Upregulated DEGs with an average log_FC value ≥ 0.5 were selected and analyzed against a reference list of all the genes detected in alignment in our experiment (n = 12,405). Statistical analysis with a Fisher’s exact test and a Bonferroni correction for multiple testing was used.

### Infection of mouse bone marrow-derived macrophages, mouse bone-marrow neutrophils and the human HL-60 cell line

C57Bl/6 mouse bone marrow-derived macrophages (BMDM) were generated as previously described [4]. Briefly, bone marrow cells were flushed from dissected femurs and tibias and cultured for 7 days in macrophage growth medium (RPMI with 10% heat-inactivated FBS (Sigma-Aldrich), 10% 3T3 m-CSF, and 1% glutamine (Thermo Fisher)) for 7 days. C57Bl/6 mouse bone-marrow neutrophils were purified from the bone marrow from femurs and tibias. Red blood cell lysis was performed at room temperature (RT) by incubation of bone marrow in 0.2% NaCl for 20 sec followed by the addition of 1 volume 1.6% NaCl. Cells were then washed in CM1 medium (RPMI with 10% Fetalclone I (Hyclone) and 2 mM EDTA), loaded on a Histopaque 1077/1119 (Sigma-Aldrich) gradient and centrifuged 30 min at 20°C. Cells at the interface of the two Histopaque solutions were collected and washed in CM1. The human HL-60 cell line (ATCC; CCL-240) was differentiated into granulocyte-like cells using 0.8% dimethylformamide (DMF) for 5-6 days.

For *in vitro* infections, bone marrow-derived macrophages and neutrophils were seeded at a density of 0.5 x 10^6^ cells/mL and 1.0 x 10^6^ cells/mL, respectively. Bacterial cultures were prepared as previously described [4]. Briefly, cells were infected with the appropriate number of bacteria to obtain a multiplicity of infection (MOI) of 1. At two hours post infection the cells were washed with the corresponding medium to remove extracellular bacteria, and remaining extracellular bacteria were killed off by incubating the cells with media containing 200 µg/mL Amikacin (Sigma-Aldrich). After 2 h the cells were washed twice with fresh medium and incubated at 32°C with 5% CO_2_ until further analysis. For analysis of intracellular growth, infected cells were lysed with 0.125% (final concentration) Triton X-100 (Thermo Fisher) for 10 min at RT. Cell lysates were 10-fold serially diluted and plated on 7H10 agar plates for CFU analysis.

### Statistical analysis

Statistical analyses were performed using the GraphPad Prism 8 software. As indicated in figure legends, a two-tailed unpaired t test or Mann-Whitney test was used for pairwise comparisons. 1-way ANOVA with Tukey’s or Kruskal Wallis comparison tests was used to compare >2 groups. For kinetic experiments or multi-parameter analysis, a 2-way ANOVA with Sidak’s or Tukey’s comparison tests was used. A p<0.05 was considered significant; *p<0.05, **p<0.01, ***p<0.001, ****p<0.0001.

## Acknowledgements

We are grateful to Christine Valfridsson, Knut Kotarsky and Marcus Svensson-Frej for technical assistance.

## Supporting Information Captions

**S1 Fig. Impact on disease, relative abundance of immune cells and on neutrophil attracting chemokine levels by *M. marinum* infection.** C57Bl/6 mice were infected with 5 x 10^6^ CFUs of WT or ΔRD1 *M. marinum*, or left uninfected (UI), as indicated. **A**) Visible tail lesions of representative mice at 28 days post infection (DPI). **B**) Cumulative length of visible tail lesions. Results (*n* = 12-13 mice) from two independent experiments. Red bars indicate the mean for each group. **C**) Bacterial burden in spleen, tail, mesenteric and tail-draining (sciatic and inguinal, pooled) lymph nodes. Results (mean + SD, *n* = 3-4 mice) representative of at least two independent experiments. 2-way ANOVA. *p<0.05, **p<0.01, ***p<0.001, ****p<0.0001. **D**, **F** and **G**) Flow cytometry analyses of infected tail tissues. **D**) Proportions of neutrophils, CD64^+^ cells and conventional dendritic cells among total CD45^+^ cells at the indicated days post infection (DPI). **E**) At the indicated DPI, tail tissues were collected and analyzed for CXCL1, 2 and 5 protein levels by ELISA. Each symbol indicates an individual mouse, and the bars show the mean for each group. Results (*n* = 8-11 infected and 7 UI mice per group; 2-way ANOVA) from two independent experiments. **F**) Proportions the G1 to G4 subpopulations among total CD64^+^ cells. **G**) Proportions of B cells, CD4^+^ and CD8^+^ T cells among total CD45^+^ cells. (**D**, **F** and **G**) Results (mean + SD, *n* = 7-9 infected or 2-3 uninfected mice per group; 2-way ANOVA). Black stars; comparison between WT and ΔRD1 infected mice. Grey stars; comparison between infected and UI mice. *p<0.05, **p<0.01, ***p<0.001, ****p<0.0001).

(TIF)

S2 **Fig. The Wasabi-encoding plasmid is stable within bacteria for at least 21 days post infection, and fluorescence correlates with bacterial colony forming units.** B6 mice were infected with 5 x 10^6^ CFUs of WT or ΔRD1 *M. marinum* carrying the Wasabi-encoding pTEC15 plasmid, or uninfected, as indicated. **A**) Representative flow cytometry plots of tail tissues analyzed for CD45 and Wasabi expression at 21 DPI. **B**) Kinetic analysis of pTEC15 plasmid (carrying a hygromycin resistance gene) curing *in vivo*. The ratio of hygromycin resistant colony forming units (Hyg^R^ CFUs) over the total amount of CFUs enumerated in the absence of hygromycin from tail samples is indicated. Results (mean + SD, *n* = 4-7 mice; 1-way ANOVA, multiple comparison to day 1 for each bacterium, ***p<0.001) from two independent experiments. **C**) Comparisons between enumerated CFUs and Wasabi^+^ cells detected by flow cytometry in the tail tissue of mice infected as indicated. Results (mean + SD, *n* = 2-4 mice).

(TIF)

**S3 Fig. *M. marinum* replicates less efficiently in neutrophils than in macrophages *in vitro*.** C57Bl/6 bone marrow-derived macrophages (BMDM), C57Bl/6 bone marrow neutrophils (BMNeu) and the human HL-60 neutrophil cell line were infected with WT *M. marinum* at MOI 1. At indicated hours post infection, intracellular growth was analyzed. Results (mean + SD, *n* = 3; 2-way ANOVA, ***p<0.001) from three independent experiments.

(TIF)

**S4 Fig. Effect of neutrophil phenotypic differentiation on expression of *Pdl1*, *Cd177* and *Hmox1*.** Single cell RNA-seq analysis of neutrophils sorted from tail tissues of mice infected with 5 x 10^7^ CFUs of WT or ΔRD1 *M. marinum* at 14 DPI. **A**) UMAP embedding of full data set (including cells from all analytical groups) with *Pdl1* or *Cd177* mRNA expression overlays. **B**) Violin plot of single cell RNA expression level within clusters. Differential expression of heme oxygenase-1 (*Hmox1*) mRNA between cluster 1 and all other clusters.

(TIF)

**S5 Fig. Role of CCR2-deficiency and iNOS activity in *M. marinum* infection.** CCR2^+/+^, CCR2^+/-^ or CCR2^-/-^ mice were infected with 5 x 10^7^ CFUs of WT *M. marinum*, as indicated. **A**) Analysis of disease development in the tail of WT infected CCR2^+/+^ and CCR2^+/-^ mice. Cumulative length of visible tail lesions was quantified at the indicated DPI. Results (*n* = 7- 14 mice per group; 2-way ANOVA). No statistical difference between the two groups. **B**) At 14 DPI, tail tissues from CCR2^+/-^ and CCR2^-/-^ infected mice were collected and analyzed for the indicated chemokines by ELISA. Results (*n* = 6 and 9 mice per group for CCR2^+/-^ and CCR2^-/-^, respectively; two-tailed Mann-Whitney test, *p<0.05, **p<0.01) from two independent experiments. **C**) B6 mice infected with 5 x 10^7^ CFUs of WT *M. marinum* were treated with aminoguanidine (AG), or PBS as a control, and tail tissues were analyzed for the indicated chemokines by ELISA at 17 DPI. Results (*n* = 7 mice per group; two-tailed Mann-Whitney test, *p<0.05, **p<0.01, ***p<0.001). **D**) Similarly infected B6 mice were treated with AG (or PBS as a control) as well as rat anti-mouse Ly6G (or isotype control) and mouse anti-rat *kappa* light chain antibodies, as indicated. Neutrophil numbers were analyzed by flow cytometry 17 DPI. Results (*n* = 9-11 mice per group; 1-way ANOVA, **p<0.01, ***p<0.001, ****p<0.0001).

(TIF)

**S1 Table. Differentially expressed genes defining neutrophil clusters *in vivo*.** Excel spreadsheet including all >0.25 log fold-change (FC) differentially expressed genes (DEGs) defining clusters 0 to 3. Sheet 1 (>0.25 log FC DEGs) includes all DEGs. Subsequent sheets include DEGs defining the individual clusters, as indicated. STRING network analysis is included in these sheets to visualize relationships between DEGs.

(XLSX)

**S2 Table. Detailed list of the materials and reagents used.**

(PDF)

